# Somatic structural variants driving distinct modes of oncogenesis in melanoma

**DOI:** 10.1101/2023.11.01.565187

**Authors:** Jake R. Conway, Riaz Gillani, Jett Crowdis, Brendan Reardon, Jihye Park, Seunghun Han, Breanna Titchen, Mouadh Benamar, Rizwan Haq, Eliezer M. Van Allen

**Affiliations:** Division of Medical Sciences, Harvard University, Boston, Massachusetts 02115, USA; Cancer Program, Broad Institute of MIT and Harvard, Cambridge, Massachusetts 02142, USA; Department of Medical Oncology, Dana-Farber Cancer Institute, Boston, Massachusetts 02215, USA; Department of Pediatrics, Harvard Medical School, Boston, MA 02215, USA; Boston Children’s Hospital, Boston, MA 02115, USA; Center for Cancer Precision Medicine, Dana-Farber Cancer Institute, Boston, MA 02215, USA

## Abstract

The diversity of structural variants (SVs) in melanoma and how they impact oncogenesis are incompletely known. We performed harmonized analysis of SVs across melanoma histological and genomic subtypes, and we identified distinct global properties between subtypes. These included the frequency and size of SVs and SV classes, their relation to chromothripsis events, and the role of topologically associated domain (TAD) boundary altering SVs on cancer-related genes. Following our prior identification of double-stranded break repair deficiency in a subset of triple wild-type cutaneous melanoma, we identified *MRE11* and *NBN* loss-of-function SVs in melanomas with this mutational signature. Experimental knockouts of *MRE11* and *NBN*, followed by olaparib cell viability assays in melanoma cells, indicated that dysregulation of each of these genes may cause sensitivity to PARPi in cutaneous melanomas. Broadly, harmonized analysis of melanoma SVs revealed distinct global genomic properties and molecular drivers, which may have biological and therapeutic impact.

**Statement of Significance:** The diversity of SVs in melanoma, and how they directly or indirectly impact oncogenesis, are incompletely known. Here we present analysis of melanoma SVs that reveal distinct global genomic properties and molecular drivers, some of which point to opportunities for further biological and therapeutic investigation.

## Introduction

Cutaneous melanoma is among the most highly mutated cancers due to the impact of UV mutagenesis leading to many C>T transitions across the genome(1,2). For this reason, molecular analyses of melanoma are often focused on somatic mutations, and increasingly so due to the association of tumor mutational burden (TMB) with response to immunotherapy(3–6). Somatic structural variant (SV) analyses of cutaneous melanoma whole genomes have been performed(2,7), with emphasis on the counts and frequency of SVs in this disease subtype. In contrast to cutaneous melanoma, acral and mucosal melanomas are associated with lower TMBs, with the majority of tumors showing no detectable effect of UV mutagenesis on their mutational spectrums. Instead, in these subtypes, comprehensive SV analysis identified higher SV burden than cutaneous melanomas and the presence of focal SVs targeting known cancer genes (e.g., *TERT*, *CDK4*, *MDM2*)(8,9). However, features relating SVs across histologic subtypes, or genomic (*BRAF*-, *(N)RAS*-, *NF1*-mutant, and triple wild-type (TWT)) subtypes of melanoma (which have been shown to have distinct secondary driver genes and pathways)(10), remain incompletely characterized.

Chromothripsis, a single complex genomic event characterized by several SVs clustered in genomic regions of oscillating copy number states across one or more chromosomes, has been systematically characterized in acral melanomas(9), and a subset of cutaneous melanomas available through the PCAWG consortium (n=106)(9,11). However, the relevance of chromothripsis between melanoma histological subtypes, and between genomic subtypes within cutaneous melanomas remains unknown. In contrast to chromothripsis characterization, the frequency and effect of SV events on topologically associated domains (TADs), which maintain the regulatory landscape of genes(12), remains unexplored across all melanoma histological subtypes. Disruption of boundaries between TADs has been shown to result in dysregulation of neighboring gene expression through a variety of mechanisms, including overexpression of oncogenes through enhancer hijacking(13) or inversions overlapping TAD boundaries placing genes near atypical regulatory elements (14,15).

Finally, a subset of cutaneous melanomas exhibit SBS COSMIC Mutational Signature 3(16). While this signature is associated with BRCA1/2 mutations and double-stranded break (DBS) repair deficiency in certain cancer types(16), it has also been shown to be associated with downregulation of *ATM* and other genes that function early in the DBS repair pathway in melanoma(10). However, no DSB repair-associated genomic features identified in cutaneous melanoma whole-exomes were significantly associated with signature 3, and the mechanism leading to the downregulation of *ATM* in the majority of these tumors remains unclear. SV analysis may enable the identification and characterization of various DSB repair mechanisms between signature 3 positive and negative tumors(7,10), beyond homologous recombination deficiency (HRD)-associated events that can be obtained through allelic copy number analysis(17,18). Taken together, we hypothesized that somatic SVs may inform: (i) molecularly defined subtype-specific modes of melanoma oncogenesis; (ii) regulatory disruption; and (iii) DNA repair defects that were not identifiable via somatic mutation analysis. Thus, we harmonized whole-genome sequencing (WGS) from 355 melanomas spanning 3 histological subtypes (acral, mucosal, cutaneous) to investigate the role of SVs in melanoma oncogenesis across these different axes.

## Results

### Cohort Overview and Subtype-Specific SV Patterns

We assembled and uniformly analyzed SVs in 355 patients with melanoma WGS (116 acral, 175 cutaneous, and 64 mucosal melanoma; Methods)(1,2,8,9). Of the cutaneous melanoma samples, 81, 55, 19, and 20 samples were *BRAF*-, *(N)RAS*-, *NF1*-, and TWT, respectively. The median sequencing coverage was 57X and 37X in tumor and matched normal samples, respectively, with no statistically significant difference in tumor sample coverage between the histologies (Wilcoxon-Mann-Whitney, p = 0.08; Supp. Figure 1). Additionally, there was no statistically significant difference in the median tumor purity between the histologies, ranging from 61% in mucosal melanomas to 66% in acral melanomas (Wilcoxon-Mann-Whitney, p = 0.37), while background ploidy in acral (3.3) and mucosal (2.9) melanomas were significantly higher than in cutaneous (2.1) melanomas (Wilcoxon-Mann-Whitney, p < 3.8 × 10^-5^). In total, our framework identified 106,032 somatic genomic rearrangements (> 30 bp; median events per tumor; acral: 81, mucosal: 64, cutaneous: 23,Methods; Figure 1A), consisting of 46,399 translocations (TRA), 25,401 deletions (DEL), 17,935 inversions (INV), and 16,297 duplications (DUP). Of the 46,399 TRA events, 13,075 (28%) were intrachromosomal while 33,324 (72%) were interchromosomal. Across acral, mucosal, and cutaneous melanomas, approximately 72.4%, 71.4%, and 70.7% of TRA events were interchromosomal, respectively.

**Figure 1:**
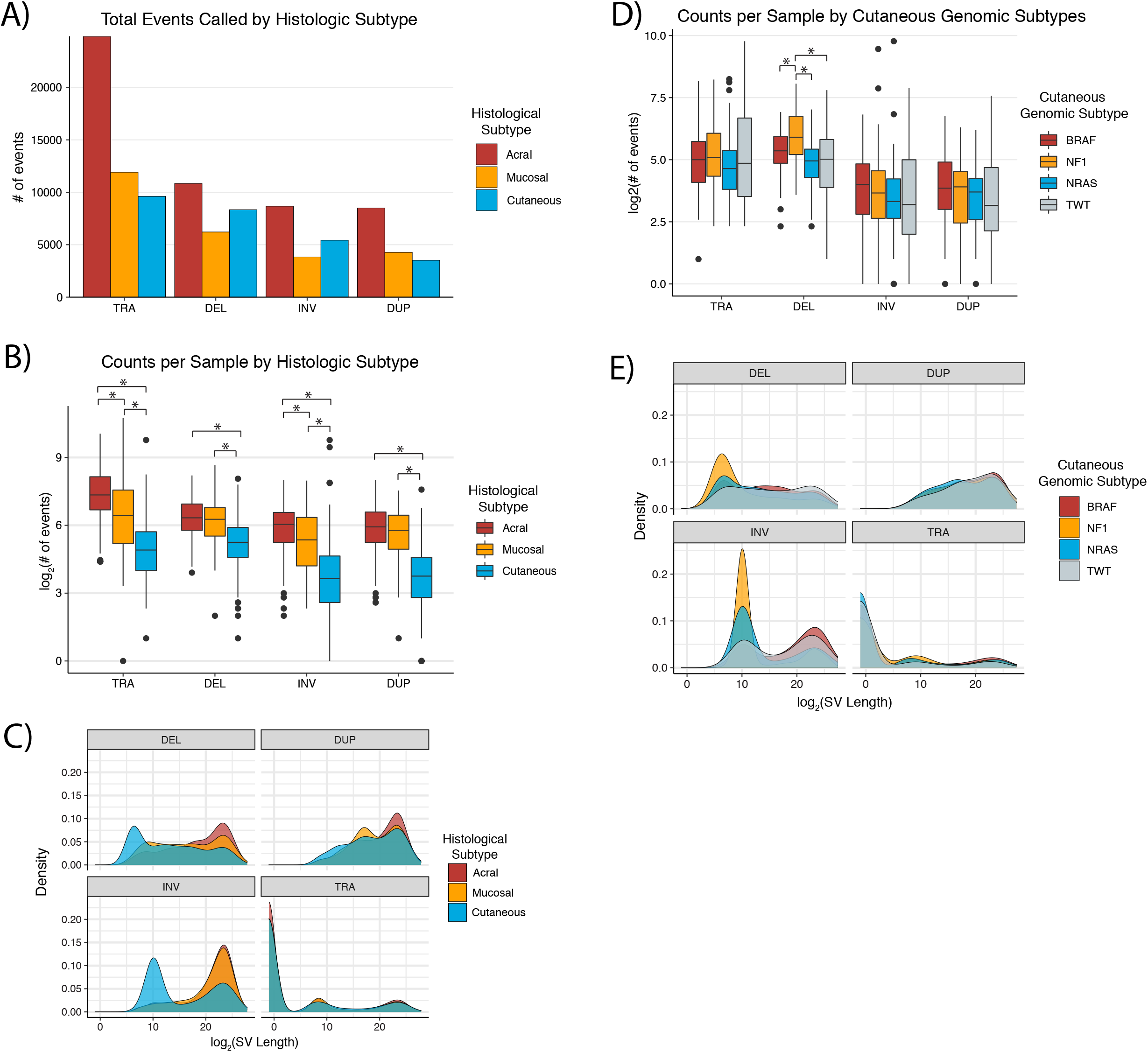
Characteristics of histologic and cutaneous genomic subtypes in melanoma. **(a)** The total number of TRA, DEL, INV, and DUP events in acral, cutaneous, and mucosal melanomas. **(b)** The distribution of the number of TRA, DEL, INV, and DUP events across acral, cutaneous, and mucosal melanoma histologic subtypes. **(c)** The distribution of the number of TRA, DEL, INV, and DUP events across the cutaneous melanoma genomic subtypes. **(d)** The distribution of the sizes of TRA, DEL, INV, and DUP events across acral, cutaneous, and mucosal melanoma histologic subtypes. **(e)** The distribution of the sizes of TRA, DEL, INV, and DUP events across the cutaneous melanoma genomic subtypes. **(b-c)** Asterisks denote a p-value < 0.05. The number of events for the **(b, d)** boxplots and SV length for the **(c, e)** density plots are plotted on a log2 scale.

The number and features of SVs varied widely across the melanoma histologies. Both acral and mucosal melanomas had significantly more events per tumor across all SV categories compared to cutaneous melanomas (Wilcoxon-Mann-Whitney, p < 2.89 × 10^-9^; Figure 1B). However, when compared to mucosal melanomas, acral melanomas had significantly higher numbers of TRA (Wilcoxon-Mann-Whitney, p = 2.9 × 10^-4^) and INV (p = 0.01) events per tumor, but not DEL or DUP events. Acral melanomas were also significantly associated with larger (measured by distance between breakpoints) SV events across all SV categories compared to cutaneous melanomas (Wilcoxon-Mann-Whitney, p < 0.026), but not mucosal melanomas. Furthermore, the distributions of DEL and INV sizes in cutaneous melanomas possessed distinctive peaks surrounding smaller SV events (< 10 kb; Kolmogorov-Smirnov, p < 2.2 × 10^-16^; Figure 1C), which may suggest a distinct mechanism of generation. Indeed, pan-cancer analysis of SVs identified small deletions to be enriched in early replicating regions near TAD boundaries, and small inversions enriched in late replicating regions(7).

Within cutaneous melanomas, there was no difference in the number of TRA, INV, and DUP events per tumor between the genomic subtypes (Wilcoxon-Mann-Whitney, p > 0.05). However, *NF1*-mutant melanomas had significantly higher numbers of DEL events per tumor compared to the other genomic subtypes (Wilcoxon-Mann-Whitney, p < 0.022; Figure 1D). Examining the distribution of DEL and INV sizes within cutaneous melanomas revealed that the majority of smaller SV events in this histology were in *NF1-* and *(N)RAS*-mutant tumors (Figure 1E). Thus, the quantity and characteristics of SVs varies widely between melanoma histologic and molecular subtypes.

### Chromothripsis and Chromoplexy Patterns in Subtypes

While chromothripsis has been identified in each of the melanoma histologic subtypes, prior studies were either unable to differentiate chromothripsis from other complex events(8,9) or were calling SVs with low sensitivity(11,19,20). Additionally, the relevance of chromothripsis between melanoma histological subtypes, or between genomic subtypes within cutaneous melanomas, is unknown (1,2,10). In this cohort, acral melanomas were significantly enriched for chromothripsis events (Methods)(11) compared to both mucosal (70% vs 31%; Fisher’s exact, OR = 5.04, 95% CI = 2.51 - 10.44, p = 8.23 × 10^-7^) and cutaneous (70% vs. 25%; Fisher’s exact, OR = 6.84, 95% CI = 3.96 - 12.04, p = 5.01 × 10^-14^; Figure 2A) melanomas, with no significant difference in the rate of chromothripsis between cutaneous and mucosal melanomas (Fisher’s exact, p = 0.41). Acral melanomas uniquely experienced more SVs than expected by chance in chromosomes 3-4, 8-10, 12-14, 16-18, 20-22, X, and Y (p < 0.05). Mucosal melanomas experienced more SVs than expected by chance in chromosome 1 (p < 0.05). Cutaneous melanomas experienced more SVs than expected by chance in chromosomes 5 (p < 0.05).

**Figure 2:**
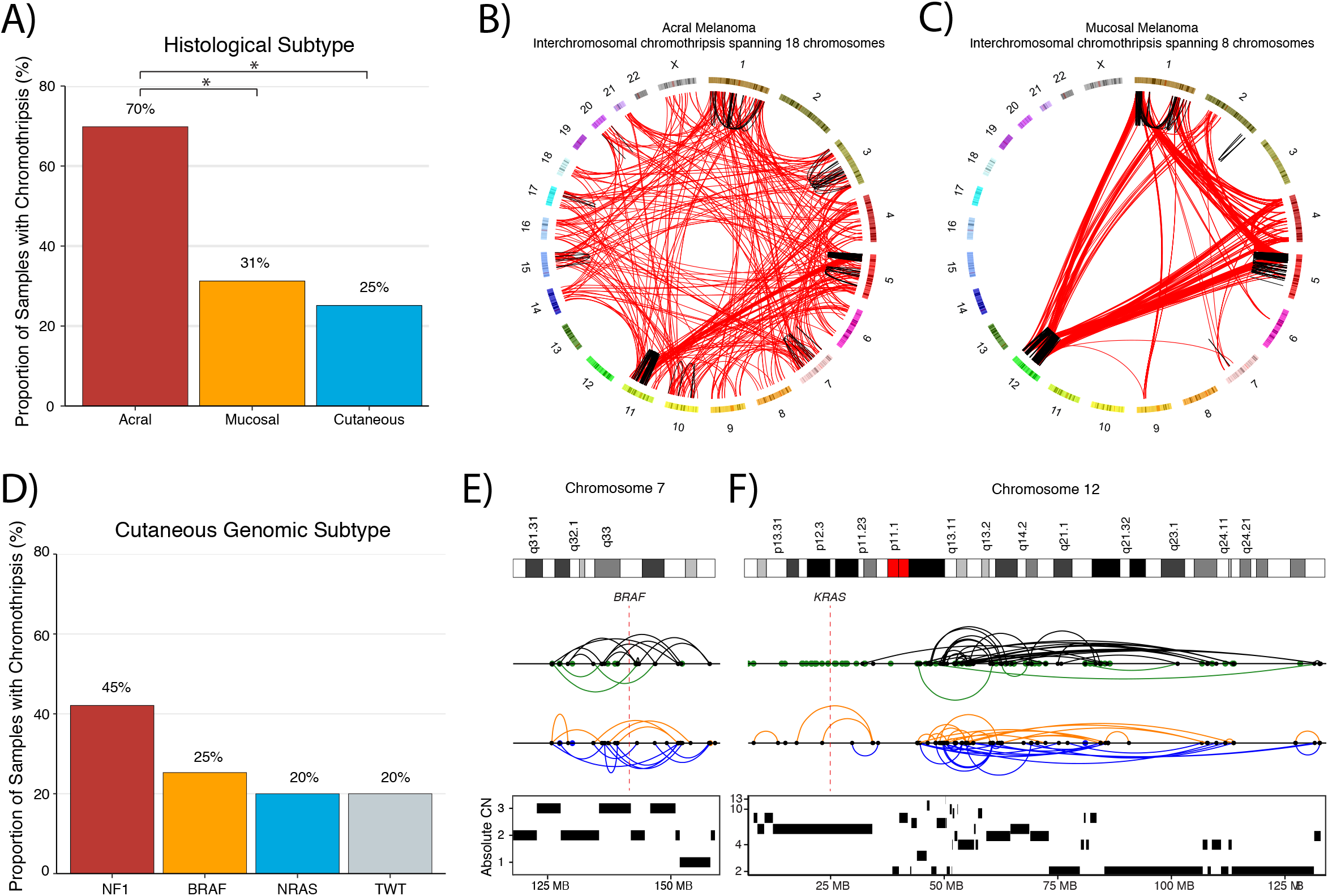
The rate and characteristics of chromothripsis events vary by melanoma histologic and cutaneous genomic subtypes. **(a)** The frequency of chromothripsis across acral, cutaneous, and mucosal melanoma histologic subtypes. **(b)** The most extreme chromothripsis event observed in an acral melanoma tumor, which consisted of SVs spanning a total of 18 chromosomes. **(c)** The most extreme chromothripsis event observed in a mucosal melanoma tumor, which consisted of SVs spanning a total of 8 chromosomes. **(d)** The frequency of chromothripsis across the cutaneous melanoma genomic subtypes. **(a, d)** Asterisks denote a p-value < 0.05. **(a, d)** Asterisks denote a p-value < 0.05. **(e)** An example of an intra-chromosomal chromothripsis event spanning the *BRAF* locus in a *BRAF*-mutant cutaneous melanoma. **(f)** An example of an intra-chromosomal chromothripsis event spanning the *KRAS* locus in a *(N)RAS*-mutant cutaneous melanoma. **(e, f)** SV events colored blue, orange, black, and green correspond to duplication-like, deletion-like, head-to-head inversions, and tail-to-tail inversions, respectively. Intrachromosomal events are connected by arches, while breakpoints of interchromosomal events are represented by points.

Approximately 85% of chromothripsis events in acral melanomas involved interchromosomal SVs, compared to 65% of mucosal (Fisher’s exact, OR = 3.05, 95% CI = 0.85 - 10.49, p = 0.055) and 52% of cutaneous (Fisher’s exact, OR = 5.17, 95% CI = 2.07 - 13.53, p = 1.1 × 10^-4^) melanomas. Of the interchromosomal chromothripsis events, the majority involved more than one additional chromosome (> 2 in total; 67% acral, 62% mucosal, and 57% cutaneous). In one extreme case, an acral melanoma tumor had a single chromothripsis event affecting 18 chromosomes (Figure 2B), whereas the greatest number of chromosomes involved in a single chromothripsis event in mucosal and cutaneous melanomas was 8 (Figure 2C) and 6, respectively. Additionally, specific chromosomes were enriched for chromothripsis in melanoma histological subtypes. Acral melanomas uniquely experienced more SVs than expected by chance in chromosomes 3-4, 8-10, 12-14, 16-18, 20-22, X, and Y (p < 0.05; Methods), mucosal melanomas experienced more SVs than expected by chance in chromosome 1 (p < 0.05; Methods), and cutaneous melanomas experienced more SVs than expected by chance in chromosomes 5 (p < 0.05; Methods). Thus, chromothripsis is associated with genomic instability in the majority of acral melanomas, while cutaneous and mucosal melanomas experience chromothripsis at less than half the rate of acral melanomas, and have similar chromothripsis landscapes despite significantly different global SV properties.

In addition to chromothripsis events, we also quantified the number of chromoplexy events. Both acral (Mann-Whitney U, p = 3.2 × 10^-4^, 3.4 vs. 2.3 events per tumor) and mucosal (Mann-Whitney U, p = 4.3 × 10^-4^, 4.2 vs. 2.3 events per tumor) melanomas were enriched for chromoplexy events compared to cutaneous melanomas. There was no difference in the number of chromoplexy events per tumor between acral and mucosal melanomas (Mann-Whitney U, p = 0.38). Within cutaneous melanomas, 42% (8/19) of *NF1*-mutant melanomas harbored chromothripsis events compared to 20-25% in the other genomic subtypes, although this did not reach statistical significance (Fisher’s exact, p = 0.09; Figure 2D). All but one (88%) *NF1*-mutant melanomas that harbored chromothripsis involved interchromosomal SVs, compared to just 38% of *BRAF*-mutant melanomas with chromothripsis. Roughly 55% and 50% of *(N)RAS*-mutant and TWT tumors with chromothripsis involved interchromosomal SVs, respectively. 2 of 3 (67%) *NF1*-mutant melanomas with missense (putatively activating) mutations in *NF1* harbored chromothripsis, compared to 6 of 16 (37.5%) *NF1*-mutant melanomas with putatively inactivating mutations in *NF1*, although this difference was not statistically significant (Fisher’s exact test, p > 0.05). There was no statistically significant difference in the proportion of V600E and V600K tumors harboring chromothripsis within *BRAF*-mutant melanomas.

A subset of samples in each genomic subtype had chromothripsis events that spanned the driver genes that define the subtypes. For example, one *BRAF*-mutant melanoma harbored an intra-chromosome chromothripsis event that affected the *BRAF* locus (Figure 2E), while 4 other *BRAF*-mutant melanomas harbored chromothripsis events that spanned (i.e. the gene is at least partially between the breakpoints of at least 1 chromothripsis generated SV) *NRAS*. One tumor with an *NRAS* G12R mutation had an intra-chromosomal chromothripsis event spanning *KRAS* (Figure 2F), while *BRAF* and *NF1* were involved in chromothripsis events in one *NRAS* melanoma each. Additionally, 2, 4, and 1 *NF1*-mutant melanomas harbored chromothripsis events spanning *BRAF*, *NRAS*, and *NF1* respectively. In TWT tumors, *BRAF* and *NRAS* were affected by chromothripsis events in one sample each. Thus, SVs generated via chromothripsis may provide secondary mechanisms of MAPK pathway dysregulation through genes that define the genomic subtypes. Furthermore, in the case of *BRAF* melanomas, these events may result in resistance mechanisms to targeted therapy(21). Thus, chromothripsis events in cutaneous melanoma may be capable of generating alterations that drive tumor initiation and development.

We lastly examined whether the distribution of short (< 10 kb) INV and DELs observed in *NF1*- and *(N)RAS*-mutant melanomas were the result of chromothripsis. The distribution of small INVs observed in *NF1*-mutant melanomas were largely driven by 2 samples, both of which had chromothripsis. However, only 34.6% and 7.3% of small INVs in these samples were located in chromothripsis regions. Similarly, the distribution of small INVs observed in *(N)RAS*- mutant melanomas was largely driven by a single sample that harbored chromothripsis, with only 8.5% of these small INVs were located in chromothripsis regions. While the distribution of short DELs observed in *NF1*- and *(N)RAS*-mutant melanomas were not driven by a few outlier samples, there again was no association with the numbers of these events and chromothripsis (Wilcoxon-Mann-Whitney, p > 0.05). These results suggest that despite the increased frequency of small SV events in *NF1-* and *(N)RAS*-mutant tumors, these events are not the result of chromothripsis, and the differences in the sizes of these SV events are driven by outlier samples.

### Effect of SVs on topologically associated domains (TADs)

Disruption of TAD boundaries through chromothripsis or other SV events can lead to the formation of neo-TADs and dysregulation of gene expression, whereby transcription factors, enhancers(13,22), and silencers(23) that are typically absent from a gene’s native TAD may act on the gene as a result of SVs(24). To investigate the effect of SVs on TADs in melanoma, we focused on SVs unlikely to span multiple TAD boundaries using an established cutoff defined by the PCAWG consortium (< 2Mb; Methods)(12). To infer the putative impact of boundary-affecting SVs (BA-SVs), we leveraged the 5 TAD type annotations from that same study(12), which were determined using the 15 chromatin state model from the Roadmap Epigenomics Project(25). These 5 TAD types are Heterochromatin, Low, Repressed, Low-Active, and Active, which are associated with increased expression (in the order specified) for genes contained within the TADs. We observed that 17.2%, 13.6%, and 7.2% of acral, mucosal, and cutaneous melanoma SVs (< 2Mb) completely spanned the full length of a TAD boundary, respectively. The frequency of these events were enriched compared to the expected number of BA-SVs based on randomly shuffled SVs, while maintaining SV size (p < 3.9 × 10^-3^; Methods). All acral melanoma tumors harbored at least one SV that spanned a TAD boundary, compared to 97% and 86.3% of mucosal and cutaneous melanomas, respectively (Figure 3A). Further, when assessing the putative functional impact of BA-SVs across histological subtypes, 97.4% of acral melanomas harbored a TAD boundary-spanning SV adjacent to an Active TAD, compared to 83% of mucosal and less than 50% of cutaneous melanomas (Figure 3B-C). While there was no significant association between chromothripsis and the presence of BA-SVs in a tumor in any histological subtype (Fisher’s, p > 0.05), tumors with chromothripsis events were associated with higher numbers of BA-SVs per tumor in acral (Wilcoxon-Mann-Whitney, p = 2.7 × 10^-5^) and cutaneous (Wilcoxon-Mann-Whitney, p = 0.026) melanomas, but not mucosal melanomas (Wilcoxon-Mann-Whitney, p = 0.09). Further, the correlation between global SV frequency and boundary altering SV frequency differed by histological subtype. The association was relatively weak in cutaneous melanomas (Pearson’s r = 0.26, p = 6.2 × 10^-4^), moderate in mucosal melanomas (Pearson’s r = 0.57, p = 8.8 × 10^-7^), and strongest in acral melanomas (Pearson’s r = 0.75, p = 2.2 × 10^-16^; Supp. Figure 2).

**Figure 3:**
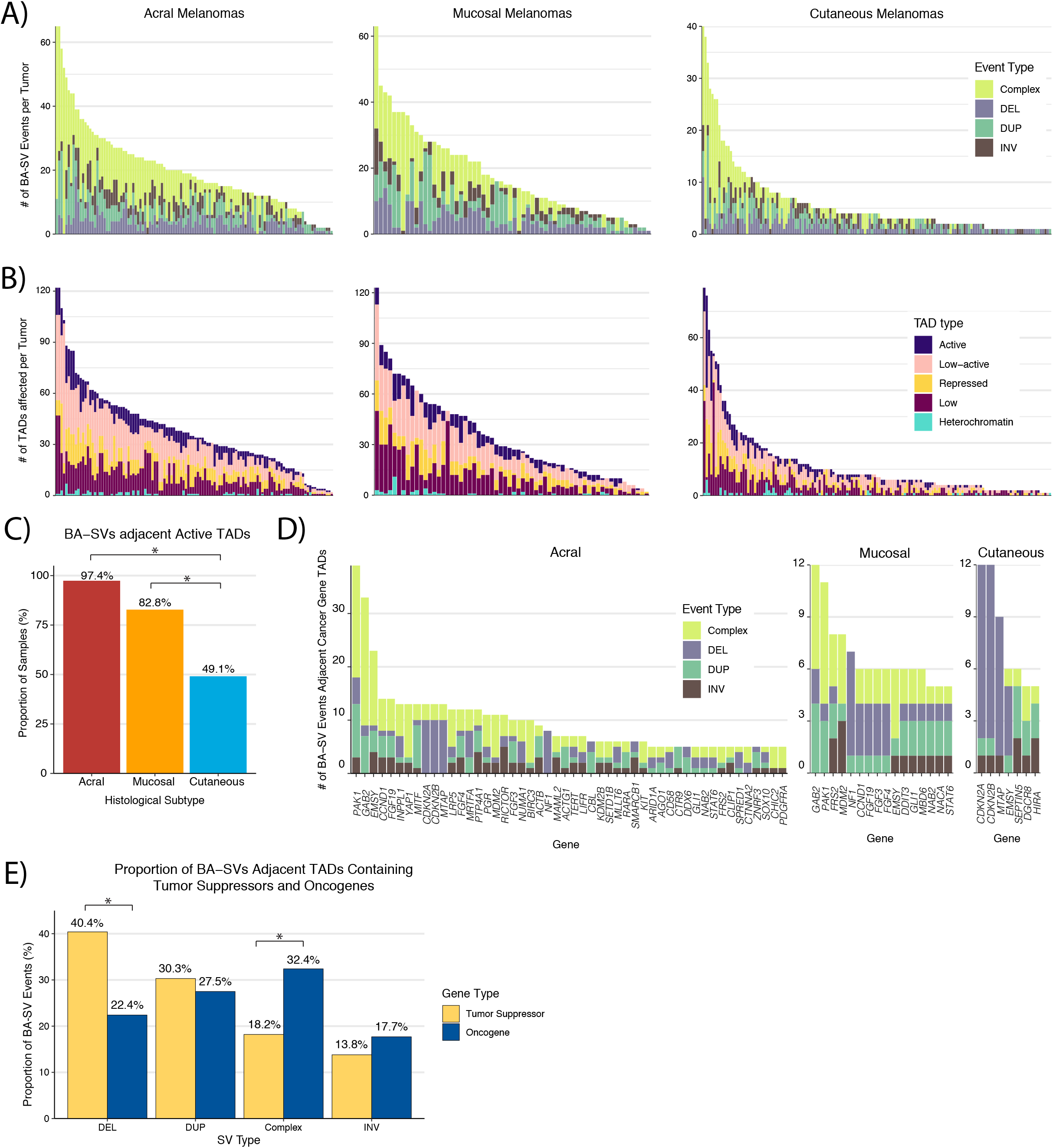
Melanomas frequently harbor SVs affecting boundaries adjacent to active TADs and TADs containing oncogenes or tumor suppressors. **(a)** The number of BA-SV spanning events per tumor across acral, mucosal, and cutaneous melanomas categorized by the type of SV event. Complex SV events are defined as overlapping concomitant DEL, DUP, INV, or TRA events. **(b)** The number of affected TADs per tumor across acral, mucosal, and cutaneous melanomas categorized by functional TAD type (Methods). **(c)** The proportion of acral, mucosal, and cutaneous melanomas with BA-SVs adjacent to active TADs. **(d)** Known oncogenes and tumor suppressors that are putatively affected by BA-SVs in at least 5 tumors per histological subtype, characterized by the type of SV event. **(e)** The proportion of event types resulting in BA-SVs that putatively affect tumor suppressors and oncogenes (Methods). **(c, e)** Asterisks denote a p-value < 0.05.

Of the total 2477 TAD boundaries, 399 (16.1%), 159 (6.4%), and 105 (4.2%) boundaries were affected by SVs in more than one tumor in the acral, mucosal, and cutaneous cohorts, respectively. Further, SVs affecting the recurrently altered boundaries (observed in at least 5 tumors) comprised 56.6%, 35.7%, and 28.4% of all boundary-spanning SVs in acral, mucosal, and cutaneous melanomas, respectively, and frequently affected TADs containing known cancer-associated genes (Figure 3D). Although we did not possess matched expression data for human samples with SVs affecting these genes (Figure 3D), orthogonal analysis in melanoma cell lines from CCLE demonstrated that SVs affecting these genes have functional consequences (Supp. Figure 3). There was no enrichment in the types of TADs adjacent to recurrently altered boundaries (altered in > 1 sample) compared to boundaries only altered in a single tumor across the histological subtypes (Fisher’s exact, p > 0.05). In general, BA-SVs adjacent TADs containing tumor suppressors (Supp. Table 1) were enriched for deletion events (Fisher’s exact; OR = 2.34; 95% CI = 1.63 - 3.35; p = 2.21 × 10^-6^; Methods; Figure 3E), whereas BA-SVs adjacent TADs containing oncogenes (Supp. Table 1) were enriched for complex events (chromothripsis or overlapping concomitant SVs; Fisher’s exact; OR = 2.62; 95% CI = 1.69 - 4.18; p = 2.71 × 10^-6^; Methods; Figure 3E).

The most recurrently affected TAD boundary in both acral (n=27, 23%) and mucosal (n=7, 11%) melanomas was chr11:77750000-77825000, which is adjacent to TADs containing the cancer genes *GAB2* and *PAK1* (Figure 4A-B). *PAK1* is an oncogene that is involved in activation of the MAPK pathway(26), and has been suggested as a potential target in *BRAF* wild-type melanomas(27). Further, *PAK1* has been identified as the most recurrently altered kinase gene via fusion events in a smaller cohort of acral melanomas(2), suggesting *PAK1* may also frequently activate the MAPK pathway outside of boundary-affecting SV events. Similarly, *GAB2* is involved in the activation of the MAPK and PI3K/AKT pathways, and has been proposed to play a role in angiogenesis in melanomas(28). This TAD boundary was altered in 4 cutaneous melanomas and was 650kb away from a fragile site (FRA11H)(29). The most recurrently altered boundary in cutaneous melanomas (all DEL events; n=7, 4%) was chr9:21700000-21775000, which is flanked by a Repressed TAD and a Low-active TAD (Figure 4C). This boundary is adjacent to the TADs containing the cancer genes *CDKN2A*, *CDKN2B*, and *MTAP*, all of which are tumor suppressors, and this boundary is located within a fragile site region (FRA9C)(30). One potential mechanism of these BA-SVs is a long range silencer interaction between regulatory elements of the adjacent repressed TAD and these tumor suppressors(31).

**Figure 4:**
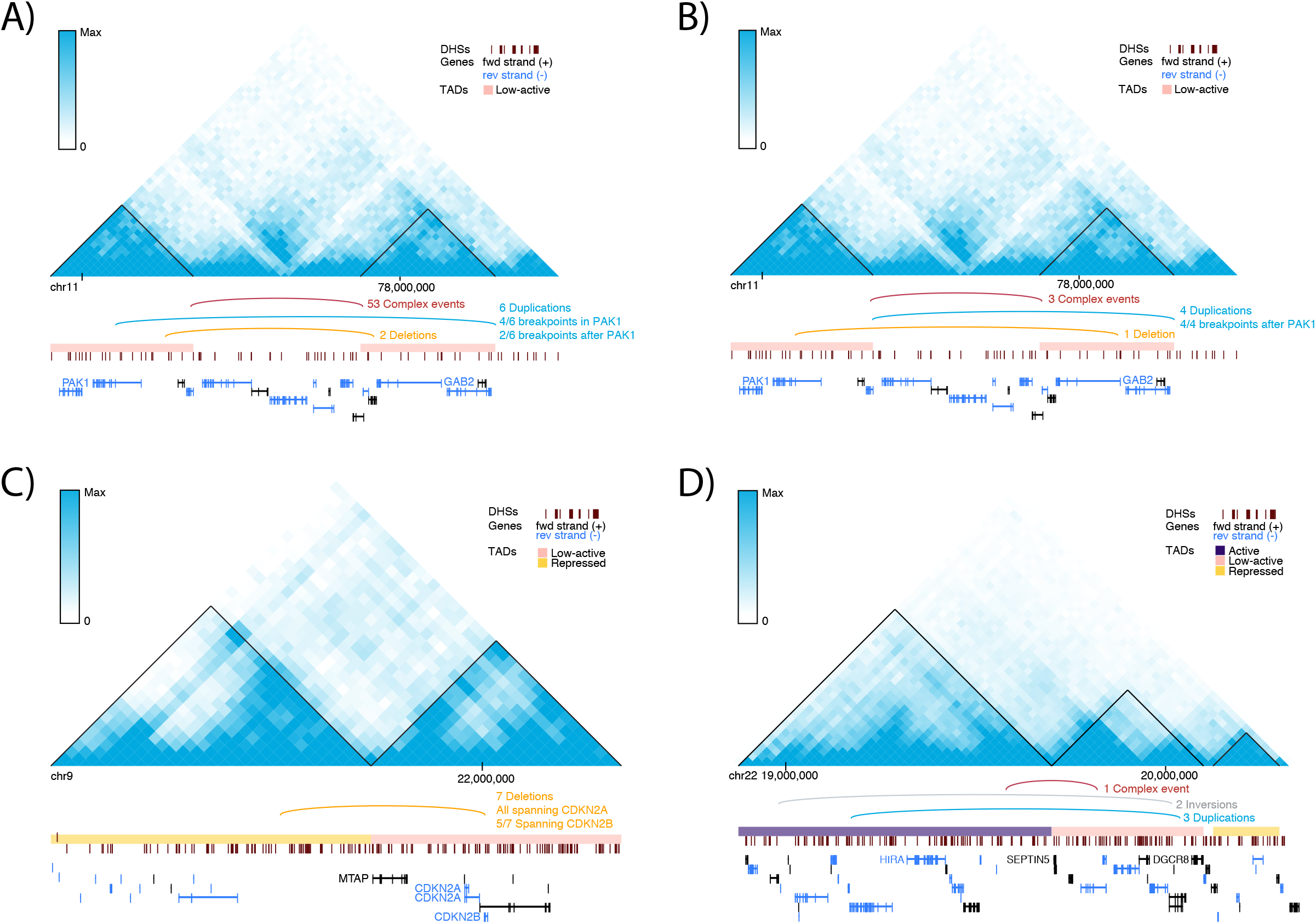
Recurrently affected boundaries adjacent to cancer gene-containing TADs. **(a)** The contact frequency map and annotations of SV events for the most recurrently altered TAD boundary in acral melanomas. **(b)** The contact frequency map and annotations of SV events for the most recurrently altered TAD boundary in mucosal melanomas. Cancer genes of interest in the adjacent TADs for both acral and mucosal melanomas include *PAK1* and *GAB2*. Both of the adjacent TADs for this boundary were low-active TADs. **(c)** The contact frequency map and annotations of SV events for the most recurrently altered TAD boundary in cutaneous melanomas. Cancer genes of interest in the adjacent TADs include *CDKN2A*, *CDKN2B*, and *MTAP*. The TAD containing these genes is a low-active TAD, and the other adjacent TAD is a repressed TAD. **(d)** The contact frequency map and annotations of SV events for the second most recurrently altered TAD boundary in cutaneous melanomas. Cancer genes of interest in the adjacent TADs include *HIRA*, *SEPTIN5*, and *DGCR8*. *HIRA* is present in an active TAD, and *SEPTIN5* and *DGCR8* are present in a low-active TAD. The contact frequencies shown here are from the IMR90 cell line, one of the 5 cell lines used to determine the functional TAD classifications by the PCAWG consortium.

The second most recurrently altered TAD boundary (chr22:19600000-19675000) was flanked by Active and Low-Active TADs (Figure 4D), and is adjacent to TADs containing the cancer genes *SEPTIN5*, *DGCR8*, and *HIRA* (Methods)(32). Unlike the other highly recurrently altered TAD boundaries, this TAD boundary was located several megabases away from the nearest fragile site (8Mb, FRA22B). Both *DGCR8* and *HIRA* are involved in UV-induced DNA damage repair, where *DGCR8* is required for transcription-coupled nucleotide excision repair (NER) at UV-induced lesions(33), and *HIRA* is a histone regulator required for efficiently priming chromatin for transcriptional reactivation following DNA repair at UV-induced lesions(34,35).

These results suggest an unappreciated role of BA-SVs in tumor development and progression across melanoma histological subtypes(12), and that BA-SVs may generate histology-enriched driver events in melanoma. Further, a subset of cutaneous melanomas experience BA-SVs affecting NER genes that may exacerbate the effect of UV mutagenesis on the mutational spectrum of tumors.

### Relationship between mutational signatures and SVs in cutaneous melanoma

To further assess the potential functional impact of SVs in melanoma, we next assessed SV pattern relationships with mutational signatures (Methods). The predominant mutational signatures in cutaneous melanoma are signature 1 (aging), signature 7 (UV mutagenesis), signature 11 (alkylating), and signature 3 (DSB repair), the lattermost of which is enriched in TWT melanomas(10). We previously reported an association between signature 3 and indel signature 8 (ID8; Non-homologous end-joining (NHEJ)), as well as between signature 3 and HRD associated copy number events, in cutaneous melanoma; however, the relationship between mutational signatures and SVs in cutaneous melanoma has remained unexplored(10). Consistent with prior analyses, mutational signature 3 was enriched in TWT cutaneous tumors in our cohort (Fisher’s exact; 5/20 vs. 5/155; OR = 9.75; 95% CI = 2.00 - 47.89; p = 1.1 × 10^-3^; Figure 5A; Methods), and it was the only SNV signature that was associated with increased numbers of SVs per tumor, after correcting for disease stage, genomic subtype, coverage, and tumor purity (multivariate regression, p = 3.2 × 10^-3^). Specifically, this association was due to increased numbers of DUP and TRA SV events (multivariate regression, p = 3.2 × 10^-4^; Figure 5B; Supp. Figure 4), but not DEL or INV SV events (multivariate regression, p > 0.17). Further, when characterizing SVs as being generated by either NHEJ, microhomology-mediated end-joining (MMEJ), or single-strand annealing (SSA), which are DSB repair mechanisms frequently involved in the repair of SV events and associated with distinct microhomology patterns at SV breakpoint junctions (Methods), signature 3 tumors were significantly associated with increased numbers of SVs arising from NHEJ (multivariate regression, p = 6.7 × 10^-3^), and decreased numbers of SVs arising from SSA (multivariate regression, p = 2.8 × 10^-4^). The ratio of NHEJ-associated SVs to SSA-associated SVs is also significantly higher in signature 3 tumors (Wilcoxon-Mann-Whitney, p = 1.95 × 10^-3^; Figure 5C). Although a smaller effect size, higher relative contribution of UV mutagenesis to the mutational spectrum of cutaneous melanomas was associated with lower numbers of SVs (multivariate regression, p < 2.7 × 10^-3^), particularly TRA and DUP events (multivariate regression, p < 4.09 × 10^-5^). There was no association between SNV mutational signatures and chromothripsis (multivariate regression, p > 0.07).

**Figure 5:**
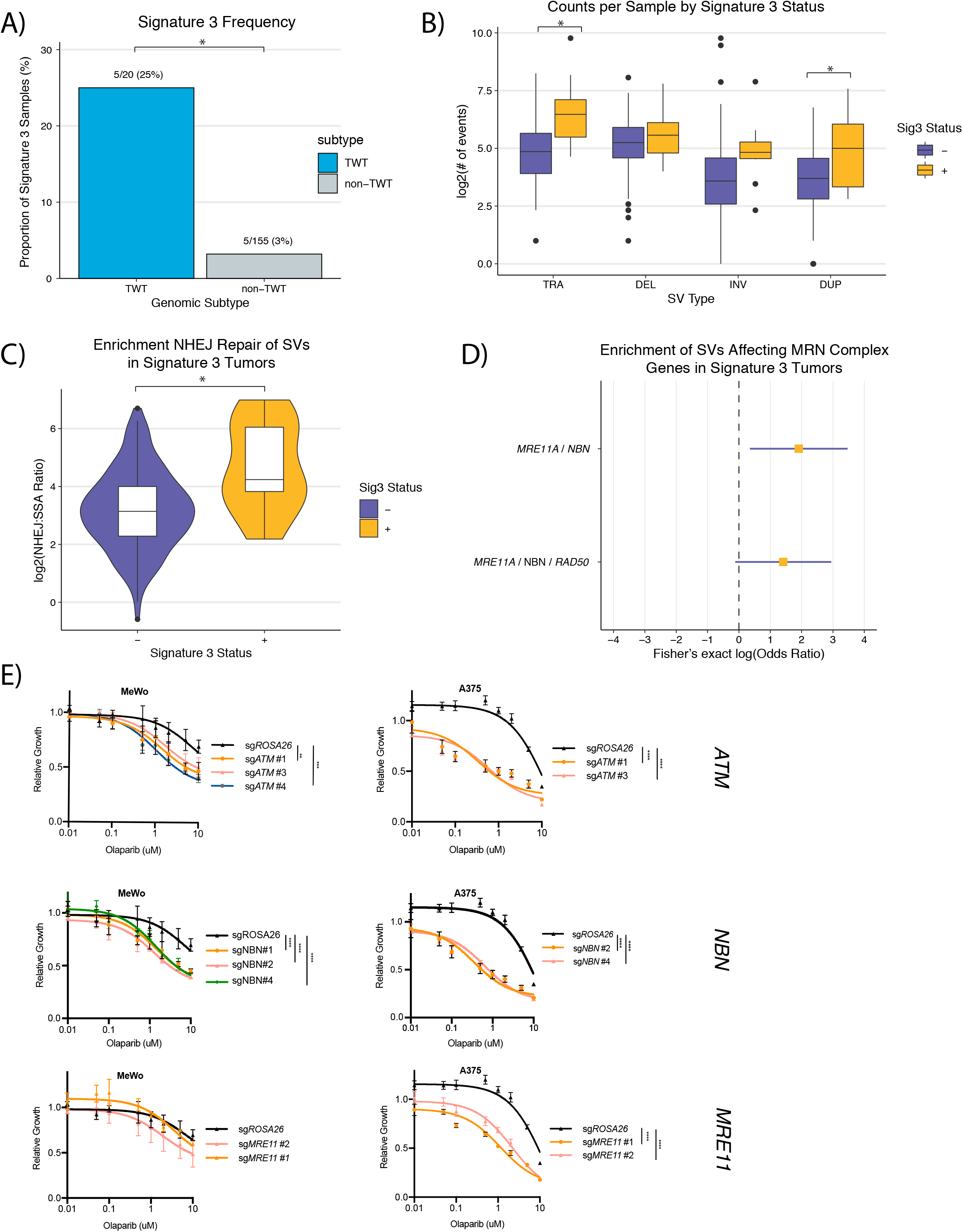
Cutaneous signature 3 melanomas are enriched for SVs frequently caused by NHEJ and are associated with SVs affecting the MRN complex. **(a)** The frequency of mutational signature 3 in TWT and non-TWT cutaneous melanomas. **(b)** The distribution of the number of events per tumor between signature 3 and non-signature 3 cutaneous melanomas, characterized by SV type. **(c)** The distribution of the ratio of putative NHEJ to SSA-generated SV events between per tumor by signature 3 status in cutaneous melanomas. **(d)** The odds ratio (yellow square) and 95% confidence interval of the odds ratio (purple line) via Fisher’s exact test for SVs overlapping MRN complex genes in signature 3 cutaneous tumors compared to non-signature 3 cutaneous tumors. **(e)** Olaparib sensitivity curves in one TWT (MeWo) and one *BRAF*-mutant (A375) melanoma cell lines with knockouts of *ATM*, *NBN*, and *MRE11*. Asterisks (*) denote adjusted p-values of: p > 0.05 (ns), p ≤ 0.05 (*), p ≤ 0.01 (**), p ≤ 0.001 (***), p ≤ 0.0001 (****) (Methods).

### SVs affecting canonical cancer genes and mutational processes in cutaneous melanoma

We then evaluated whether specific SVs affected canonical cancer genes and may directly relate to the mutational processes observed in cutaneous melanoma. Similar to our finding that cutaneous melanomas possessed somatic mutations in distinct secondary driver genes(10), several canonical cancer genes were also enriched for SVs within each genomic subtype. The most significantly enriched alterations in *BRAF*-mutant compared to non-*BRAF*-mutant melanomas were non-duplication SV events in *CDKN2A* (39/81, 48%; Fisher’s exact, OR = 2.41, 95% CI = 1.24 - 4.78, p = 7.8 × 10^-3^). Only 1 *BRAF*-mutant and 1 *BRAF*-wild-type melanoma had duplication events overlapping *CDKN2A*. *NF1*-mutant melanomas were significantly associated with non-duplication SV events in two RASopathy genes, *RAF1* and *SPRED1* (Fisher’s exact, OR = 5.03, 95% CI = 1.46 - 16.42, p = 4.8 × 10^-3^), the latter of which has also been identified as a significantly mutated gene exclusive to *NF1*-mutant melanomas(10,36). The most statistically significant canonical cancer gene affected by SVs in TWT melanomas was *CBFA2T3*, which was not altered in any of the other genomic subtypes (Fisher’s exact, OR = Inf, 95% CI = 5.69 - Inf, p = 1.3 × 10^-4^), and is a putative tumor suppressor in breast cancer(37,38). *CBFA2T3* exclusively harbored TRA and INV events in TWT tumors (n=4).

*MRE11* was also among the cancer genes significantly enriched for SVs in TWT tumors compared to other subtypes (Fisher’s exact, OR = 5.36, 95% CI = 1.01 25.18, p = 0.024), and is one of the core genes of the MRN complex, along with *NBN* and *RAD50*, and is involved in the initial processes of DSB repair prior to homologous recombination and NHEJ, and is responsible for activating *ATM*(*39,40*). We previously found that signature 3 in TWT tumors was associated with downregulation of *ATM*, although we were unable to identify recurrent alterations in somatic coding regions that might explain the downregulation of *ATM* in a subset of samples(10). Three of the 5 TWT tumors with SVs affecting *MRE11* had signature 3. Expanding the analysis to all signature 3 vs. non-signature 3 tumors also revealed the enrichment of *NBN* in signature 3 tumors (Fisher’s exact, OR = 7.26, 95% CI = 1.04 - 39.44, p = 0.023), another core gene of the MRN complex. All SVs affecting *MRE11* and *NBN* in signature 3 tumors were complex events (Methods), compared to less than half (43%) of non-signature 3 tumors (Fisher’s exact, OR = 6.74, 95% CI = 1.42 - 32.04, p = 7.4 × 10^-3^; Figure 5D). Pathway overrepresentation analysis (Methods) on the set of cancer genes significantly enriched for SVs in signature 3 tumors compared to others identified the MRN complex as the top enriched protein complex (q = 1.79 × 10^-3^).

Although SVs affecting *RAD50* were not associated with signature 3 tumors, there was no difference in the association of *MRE11* (r=0.73) or *NBN* (r=0.72) expression with *ATM* expression compared to the association of *RAD50* expression (r=0.73) with *ATM* expression in TWT tumors (Supp. Figure 5A). However, the correlation between MRN complex expression and *ATM* expression was significantly stronger in TWT tumors than in non-TWT tumors (r=0.82 vs r=0.69; Fisher’s Z-transformation, p = 0.03; Supp. Figure 5B). To assess whether the correlation observed in TWT tumors was not spurious due to having 7-fold fewer samples, we performed downsampling analysis for 10,000 simulations (Methods). Only 2.57% of these downsampled simulations yielded a correlation coefficient higher than that initially observed for TWT tumors (p = 0.0257, Move to Supp. Figure 5C). These results suggest that MRN-dependent *ATM* activation may be more frequent in TWT tumors or that *ATM* activation is more tightly regulated by the MRN complex in TWT tumors, potentially explaining why the association between signature 3 and *ATM* downregulation was restricted to TWT tumors. Additionally, these results are consistent with our previous finding that signature 3 in TWT cutaneous melanomas are associated with dysregulation of *ATM* and affects genes that function early during the initiation process of DSB repair.

To determine if melanomas that have dysregulation of the MRN complex or *ATM* may be sensitive to PARP inhibitors (PARPi), we performed independent knockouts of *ATM*, *NBN*, and *MRE11* in 2 melanoma cell lines, MeWo (TWT) and A375 (*BRAF*-mutant), followed by olaparib cell viability assays (Methods; Supp. Figure 6). MeWo cells lacking *ATM* and *NBN*, but not *MRE11*, showed increased sensitivity to Olaparib (Figure 5E), and A375 cell lines lacking *ATM*, *NBN*, and *MRE11* all showed increased sensitivity to olaparib (Figure 5E). These results suggest that while dysregulation of *ATM* and the MRN complex is specifically enriched in TWT melanomas, this dysregulation may be sufficient to cause DSB repair deficiency in both TWT and non-TWT tumors, possibly rendering them sensitive to PARPi.

## Discussion

Through uniform analysis of SVs on the largest melanoma WGS cohort to date, we revealed distinct frequencies and putative drivers of melanoma histological and cutaneous genomic subtypes. Acral and mucosal melanomas were associated with more SVs per tumor relative to cutaneous melanomas regardless of the SV type, and acral melanomas were enriched for chromothripsis events relative to cutaneous and mucosal melanomas. Additionally, in tumors that had chromothripsis events, acral melanomas were associated with higher rates of interchromosomal chromothripsis events compared to cutaneous and mucosal melanomas. While the frequencies of SV events differed between acral and mucosal melanomas, the functional impact and driver gene alterations observed in these histological subtypes were similar. Roughly 97% and 83% of acral and mucosal melanomas, respectively, had BA-SVs that affected functionally active TADs compared to less than half of cutaneous melanomas. In cutaneous melanomas, *NF1*-mutant tumors were enriched for deletion SVs, and had chromothripsis events at nearly twice the rate compared to the other genomic subtypes.In addition to having the highest TMB of the genomic subtypes, *NF1*-mutant tumors also have the highest SV burden(1,10). While our study emphasized assessments of subtype-specific SV processes for translational purposes (i.e. DNA repair deficiencies and therapeutics), exploration of additional complex events such as tyfonas, rigmas, and pyrgos may prove to be clinically and functionally relevant(41). Additionally, analysis of double-minute events through assays such as FISH(42), as well as understanding the timing of molecular events with spatial or temporal samples (i.e. via TRACERx(43)), may guide further interpretation of SVs in melanoma.

BA-SVs can disrupt the 3-dimensional architecture of the genome, and expose genes to sets of functional elements they normally wouldn’t interact with, thereby resulting in dysregulation of the genes(13)(14,15). Of the 16 genes recurrently affected (observed in at least 5 tumors) by BA-SVs in mucosal melanomas, 13 were shared with acral melanoma. One of these genes was *NF1*, one of the MAPK pathway genes used to define the cutaneous melanoma genomic subtypes. All but one BA-SV affecting *NF1* in acral and mucosal melanomas were deletion events. In addition to sharing similar recurrently affected genes, acral and mucosal melanomas also shared the the most recurrently altered TAD boundary (chr11:77750000-77825000), which is adjacent to the TADs containing *PAK1* and *GAB2*. Only *EMSY* was recurrently affected by BA-SVs in both cutaneous and mucosal melanomas, and only *CDKN2A* and *CDKN2B* were recurrently affected by BA-SVs in both acral and cutaneous melanomas. Thus, despite having drastically different SV landscapes, acral and mucosal melanomas share many of the same putative driver SVs. This is akin to the shared somatic mutation-derived driver genes from these subtypes(2,44).

The most recurrently affected genes by BA-SVs in cutaneous melanomas were *CDKN2A* and *CDKN2B*, with the majority being deletion events. *CDKN2A* was also enriched for non-duplication events (39/81; 48%) in *BRAF*-mutant cutaneous melanomas compared to the other genomic subtypes. Notably, *CDKN2A* has also been identified as a canonical driver in cutaneous melanoma via analysis of somatic mutations(1,10). The second most recurrently altered TAD boundary in our cutaneous melanoma cohort affected the NER genes *HIRA* and *DGCR8*, which are involved in the repair of mutations caused by UV mutagenesis. Increased activity of UV mutagenesis is associated with higher TMB(45), and therefore may have implications for immunotherapy treatment decisions or response. Other than *MTAP* and *SEPTIN5*, *CDKN2A*, *CDKN2B*, *HIRA*, and *DGCR8* were the only recurrently altered cutaneous melanoma genes that were not recurrently altered in acral or mucosal melanomas, which frequently lack the presence of UV-induced mutations.

Mutational significance analysis in cutaneous melanoma has revealed that the genomic subtypes preferentially experience mutations that affect distinct pathways. *NF1*-mutant melanomas preferentially harbored alterations in RASopathy genes, with *SPRED1*, *RASA2*, and *RASSF2* being identified as significantly mutated genes within the subtype(10). *NF1*-mutant melanomas in our cohort were enriched for SV events affecting *SPRED1* and *RAF1*, the latter of which has been implicated in activating fusion events in cutaneous melanoma and enriched in TWT tumors(46).

A subset of cutaneous melanoma tumors have been characterized as having mutational signature 3 (associated with DSB repair deficiency), enriched in TWT tumors(10). While the prevalence of signature 3 has been characterized in cutaneous melanoma WGS samples, its association with SVs has remained unexplored. Here we show that signature 3 is associated with increased DUP and TRA SV events in melanoma, and is associated with a higher rate of the error-prone NHEJ repair. However, we did not find a significant association between signature 3 and chromothripsis, despite prior studies linking HRD to increased prevalence of chromothripsis(47). Signature 3 in TWT melanoma tumors is associated with downregulation of *ATM* and methylation of *INO80*, however the source of *ATM* downregulation is unknown. Here we identified the enrichment of SVs affecting MRN complex genes in signature 3 tumors, which directly interacts with *ATM*. Like *ATM* and *INO80*, the MRN complex functions early in the DSB repair pathway, providing further evidence for the source of signature 3. Cell viability assays in TWT and non-TWT cell lines with either *ATM*, *NBN,* or *MRE11* knocked out revealed that melanoma cell lines lacking the expression of these genes are sensitive to Olaparib, and warrants a more exhaustive follow-up to determine if PARPi may benefit molecularly stratified melanoma patients in the clinic.

Overall we demonstrated that SV analysis of melanoma whole-genomes can identify additional putative driver mechanisms unique to histological and cutaneous genomic subtypes, some of which may present as clinically relevant druggable events. Still, further experimental work in preclinical models will be required to determine the therapeutic relevance of MRN complex alterations and *ATM* downregulation in melanoma, as well as the functional consequences of SVs at recurrently altered TAD boundaries. Furthermore, the number of whole-exome samples far exceeds the number of whole-genome samples in melanoma. Continued harmonized molecular analysis of a larger melanoma WGS cohorts will help determine the robustness and true prevalence of potential driver alterations identified in this study.

## Methods

### Whole-genome sequencing dataset description

We downloaded publicly available aligned WGS BAM files from 4 previously published studies (Supp. Table 2). For SV analysis, we required both tumor and normal samples to have a sequence coverage of at least 20X, and a tumor purity of at least 20%. The median sequencing coverage was 57X in the tumor samples and 37X in the normal samples. The median tumor purity ranged from 61% in mucosal melanomas to 66% in acral melanomas.

The cutaneous melanoma mutation data, which was used to determine genomic subtype and identify mutational signatures (see Mutational signatures), was downloaded from the supplement of Hayward *et al*. 2017(2) and the ICGC Data Portal (https://dcc.icgc.org/)(48) for TCGA-SKCM WGS samples.

### TCGA RNA-seq Data

The cutaneous melanoma expression data used in this study is from the TCGA-SKCM cohort, which is publicly available from the TCGA-SKCM workspace on Terra (TCGA_SKCM_ControlledAccess_V1-0_DATA) via dbGaP access. The RSEM upper quartile normalized bulk RNA expression data was used for all expression analysis in this study, and can be found under the following column identifier in the Terra workspace: rnaseqv2 illuminahiseq_rnaseqv2 unc_edu Level_3 RSEM_genes_normalized data.

### SV calling

We called SVs with three different SV calling methods: Manta (https://github.com/Illumina/manta)(49), DELLY2 (https://github.com/dellytools/delly)(50), and SvABA (https://github.com/walaj/svaba)(51). To identify a set of high-confidence SVs per tumor we filtered the calls to only keep SVs identified by 2 or more methods, in accordance with best practices(52), allowing for a maximum distance of 1kb pairwise between breakpoints and requiring that the calls agree on type, strand, and are at least 30bp long. This filtering was performed using the SURVIVOR R package (https://github.com/fritzsedlazeck/SURVIVOR)(53).

### SV annotations

To add gene-level annotations to our high confidence SV set, we ran AnnotSV v3.0(https://lbgi.fr/AnnotSV/)(54) using the default set of hyperparameters. The SV annotations were run on December 29th, 2020.

### Copy number calling

Allelic copy number calls were determined using FACETS (https://github.com/mskcc/facets)(17), which also provides tumor purity and ploidy information. These copy number calls were used as input to ShatterSeek(11) (see Identification of chromothripsis events) for identifying the oscillating copy number criteria of chromothripsis events.

### Identification and visualization of chromothripsis events

To identify chromothripsis events in melanoma cancer genomes we ran ShatterSeek (https://github.com/parklab/ShatterSeek)(11) using the high-confidence SVs and allelic copy number data from FACETS as input. ShatterSeek was also used to visualize chromothripsis events on single chromosomes, such as in Figures 2E-F. To visualize interchromosomal chromothripsis events we used the circos tool on Galaxy (https://usegalaxy.org/)(55). Non-chromothripsis complex SV events were defined as overlapping concomitant DEL, DUP, INV, or TRA events that were not determined to be caused by chromothripsis via Shatterseek. This is the same definition of non-chromothripsis complex SV events as Akdemir *et al.*, 2020(12) (PCAWG), which classifies these as SV events that involve multiple junctions, but do not include SVs that are involved in chromothripsis events as determined by ShatterSeek.

### Chromothripsis chromosomal enrichment

To test if certain chromosomes were enriched for chromothripsis events, we randomly shuffled around the associated SV breakpoints, taking into account chromosome size. P-values were calculated as the probability that shuffling the breakpoints led to the same or more chromothripsis related SVs than observed on a given chromosome.

### TAD and TAD boundary assignments/annotations

TAD and TAD boundary assignments, as well as TAD type annotations were downloaded from Akdemir *et al*. 2020(12). Here, TAD and TAD boundary coordinate assignments were determined by identifying TAD boundaries that were within 50kb of each other across Hi-C data from 5 different cell line types (GM12878, HUVEC, IMR90, HMEC and NHEK). TAD type annotations (Heterochromatin, Low, Repressed, Low-Active, and Active) were determined by k-means clustering according to the 15 state ChromHMM model from the Roadmap Epigenomics Project(25), and associating the clusters with gene expression data from GTEx(56) and ICGC(48).

Short range SVs likely to only affect a single TAD boundary were classified as < 2Mb in length, and were the only types of SVs used in the boundary-affecting analysis. The cutoff of < 2Mb was defined by the PCAWG consortium (< 2Mb)(12). For a short range SV to be considered boundary affecting, the entire TAD boundary had to be overlapped by the SV.

### BA-SV Permutation

To determine if the frequency of BA-SVs observed in each melanoma histological subtype were enriched beyond what was expected, we performed 1000 permutations where the SVs were shuffled, while maintaining size, and overlaps with TAD boundaries were observed. The p-value was determined by the proportion of permutations that resulted in a higher frequency than was observed by the SV calls.

### Fragile site annotations

Fragile site annotations were obtained from https://webs.iiitd.edu.in/raghava/humcfs/(57). Specifically, we used the “Fragile site bed files” reference, which provides a directory of bed files containing fragile site regions on a per chromosome basis.

### Classification of double-stranded break repair mechanisms

To classify SVs as being repaired by NHEJ, MMEJ, or SSA, we applied the breakpoint microhomology cutoffs identified in Li *et al*. 2020(7), which were determined by fitting linear functions to breakpoint microhomology data across PCAWG. This resulted in the identification of 3 sets of structural variants defined by microhomologies of 1 bp, 2-9 bp, and 10 or more bp, which were classified as NHEJ, MMEJ, and SSA, respectively.

### Mutational signatures

To identify mutational signatures present in tumor samples we ran deconstructSigs (https://github.com/raerose01/deconstructSigs)(58) using the COSMIC v2 signatures reference(59,60) and a signature contribution cutoff of 0.06. This contribution cutoff provides a false-positive rate of 0.1% and false-negative rate of 1.4%, and is the recommended cutoff.

### Pathway over-representation analysis

We performed pathway over-representation analysis on the set of cancer genes enriched in signature 3 tumors via Fisher’s exact method using ConsensusPathDB (v. 34) (http://cpdb.molgen.mpg.de)(61). We ran ConsensusPathDB (on May 18th, 2021) using the default parameters for both pathway-based gene sets and protein complex-based gene sets.

### Expression correlation analysis

We performed correlation between *ATM* expression and MRN complex expression using the TCGA-SKCM RSEM upper quartile normalized bulk RNA-seq data. To calculate an aggregate expression for the entire MRN complex we calculated the geometric mean of *MRE11*, *NBN*, and *RAD50*. Correlation was calculated using the stats R package, and the geometric mean was calculated using the psych R package.

### CCLE SV functional consequence analysis

The Q2 2023 CCLE datasets were subset to melanoma cell lines only for the analysis. For each gene included in the analysis (*MITF*, *TERT*, *MAML2*), TPM expression was z-score normalized. Since only a small number of samples (*MITF* n=2, *TERT* n=1, *MAML2* n=1) had SVs affecting genes highlighted in Figure 3D, we reported the expression percentiles for each individual gene. To further determine an association between SV events and expression changes, we performed a Mann-Whitney test on the pan-gene aggregated z-scored (i.e. each gene is z-scored separately to put the expression on the same scale) expression.

### Gene sets

The oncogene and tumor suppressor gene sets used in the BA-SV analysis were downloaded from MSigDB(62,63) on May 27th, 2021 under the curated Gene Families (https://www.gsea-msigdb.org/gsea/msigdb/gene_families.jsp). The set of cancer genes were determined by taking the union of Cancer Gene Census (v. 86) genes and OncoKB(64) cancer genes.

### Statistics and reproducibility

Statistical analyses were performed using the stats R package for R v.3.6.1. Reported q-values represent FDR-corrected p-values and reported p-values represent nominal p-values. All statistical tests performed (e.g., Wilcoxon-Mann-Whitney, Kolmogorov–Smirnov, Fisher’s exact test) were two sided.

### Cell Lines

MeWo (ATCC) were cultured in EMEM supplemented with 10% FBS and 1% penicillin-streptomycin. CHL1 and A375 (ATCC) were cultured in DMEM supplemented with 10% FBS and 1% penicillin-streptomycin. Cells were incubated at 37°C in 5% CO2. Cells were tested for mycoplasma using PCR-based screening (PCR Mycoplasma Detection Kit, Cat# G238, Applied Biological Materials Inc.) weekly or biweekly.

### Cell line authentication

Cell line authentication was performed using the ATCC Sample Collection Kit Cell Authentication Service.

### CRISPR/Cas9 targeting for the generation of knockout cell lines

*ATM*, *MRE11*, *NBN* knockouts were generated in MeWo and A375 cell lines using CRISPR/Cas9-based gene editing. The sgRNAs oligos purchased from Eton Biosciences (San Diego, CA) (Supp.Table 3), were annealed and cloned into lentiCRISPR v2 (#52961) purchased from Addgene (Watertown, MA) according to published protocols(65,66). sgROSA26 was used as a negative control. The generated lentiviral plasmids were cotransfected with viral packaging plasmids PAX2 and pMD2.G into HEK 293T Lenti-X cells (Clontech) using TransIT-LT1 (Mirus Bio LLC, Madison WI). MeWo, CHL1, and A375 cell lines were infected with the lentivirus followed by drug selection using Puromycin (1ug/mL, InVivoGen). An early passage (2 weeks post viral infection) pooled puromycin-resistant population was used for the subsequent western blots and cell viability assays.

### Western Blot Analysis

Whole-cell lysates were prepared in RIPA lysis buffer (BP-115, Boston Bioproducts, Inc, Ashland MA) supplemented with cOmplete Mini protease inhibitor (Roche) and Phospho-STOP phosphatase inhibitor (Roche). BCA Protein Assay Kit (Thermo Scientific) was used to normalize protein quantities. Samples were denatured with SDS loading dye at 95 °C for 10 min. Samples were resolved on 4-20% Criterion TGX Stain-Free Precast Gels (#5678094, BioRad) at 100 V for 15 min. then 160 V for 45 min. Proteins were transferred to 0.2um TransBlot Turbo Midi size nitrocellulose membrane using Trans-Blot Turbo (BioRad). The membranes were blocked for one hour in 5% milk in TBST, washed for 10 min. in TBS-Tween then incubated with primary antibodies in 5% milk in TBST at 4 °C overnight. The antibodies used were: anti-Atm (#2873T, Cell Signaling), anti-Mre11 (#4895, Cell Signaling), anti-Nbs1 (#NB100-143SS, Novus Biologicals), and anti-Vinculin (ab129002, Abcam). After three 15 min. washes with TBST, membranes were incubated with the secondary antibody (in 5% milk in TBST) for 1 hour at room temperature. After three washes with TBST, chemiluminescence reaction was performed using ECL Western Blotting Substrate (Pierce). Films were developed in a darkroom using Kodak X-OMAT 2000A processor.

### Cell Viability Assays

Each cell line was seeded in 96-well plates (500 cells per well, CellTreat) and incubated overnight in the appropriate growth media. Olaparib (AZD2281 from Selleck Chemicals) was diluted in DMSO and added at the indicated concentrations in triplicates. After 4 days, cell viability was assessed using CellTiter-Glo Luminescent Cell Viability Assay (#G7572, Promega) and luminescence values were obtained using Infinite M200Pro plate reader (Tecan). Results were analyzed and survival curves were generated using Prism 7. Statistical analysis was performed using one-way ANOVA Dunnett’s multiple comparisons test (comparing the AUC of each curve to the control sgROSA26).

### Data Availability Statement

All WGS BAM files used in this study were accessed from publicly available cohorts, and details on cohorts for access are summarized in Supplementary Table 2. The cutaneous melanoma mutation data, which was used to determine genomic subtype and identify mutational signatures (see Mutational signatures), was downloaded from the supplement of Hayward *et al*. 2017(2) and the ICGC Data Portal (https://dcc.icgc.org/)(48) for TCGA-SKCM WGS samples.

---

**Supplementary Figure 1:**
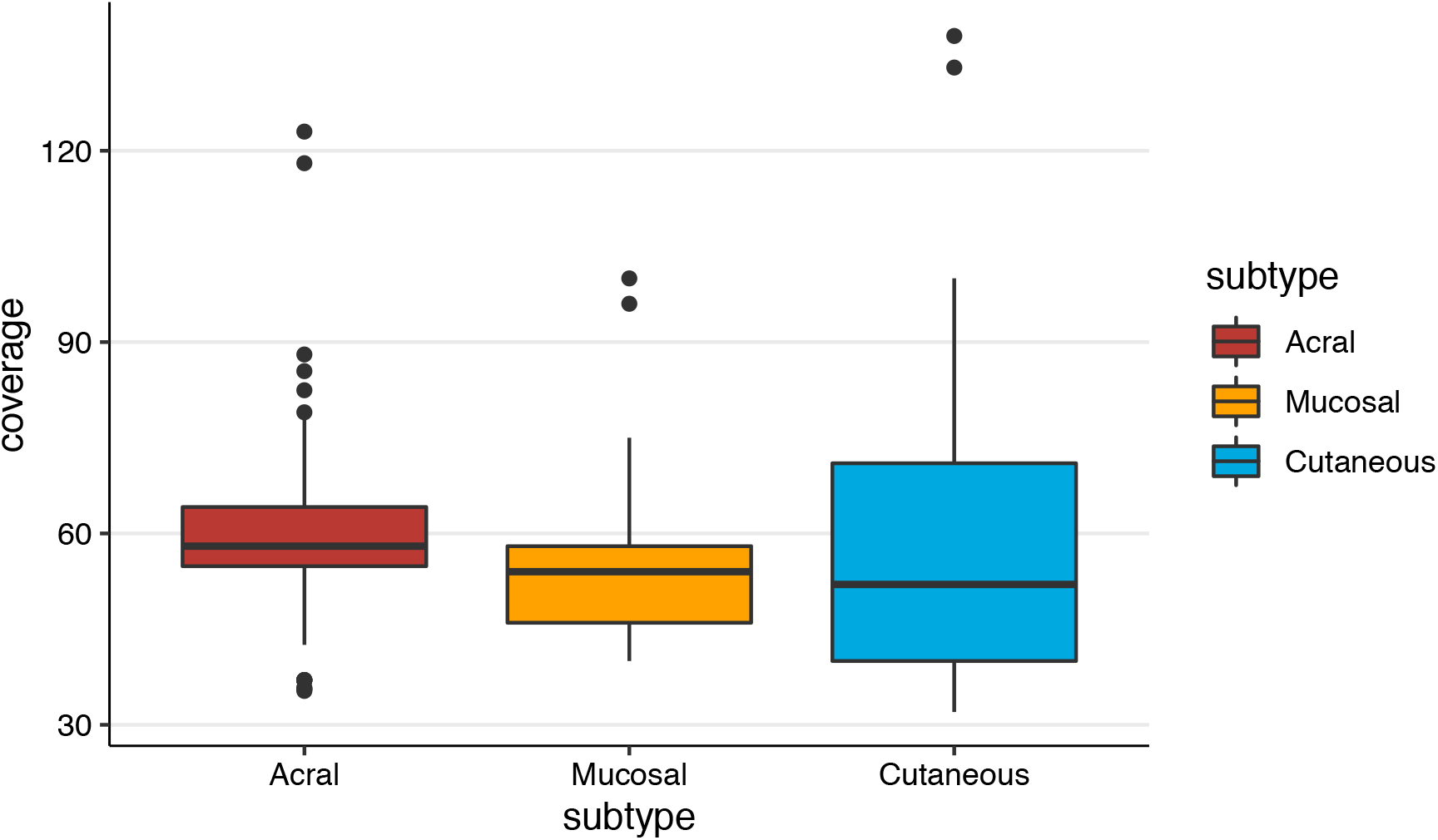
Distribution of sequencing coverage by histological subtype. There was no difference in the sequencing coverage between samples from different histological subtypes (Wilcoxon-Mann-Whitney, p = 0.08).

**Supplementary Figure 2:**
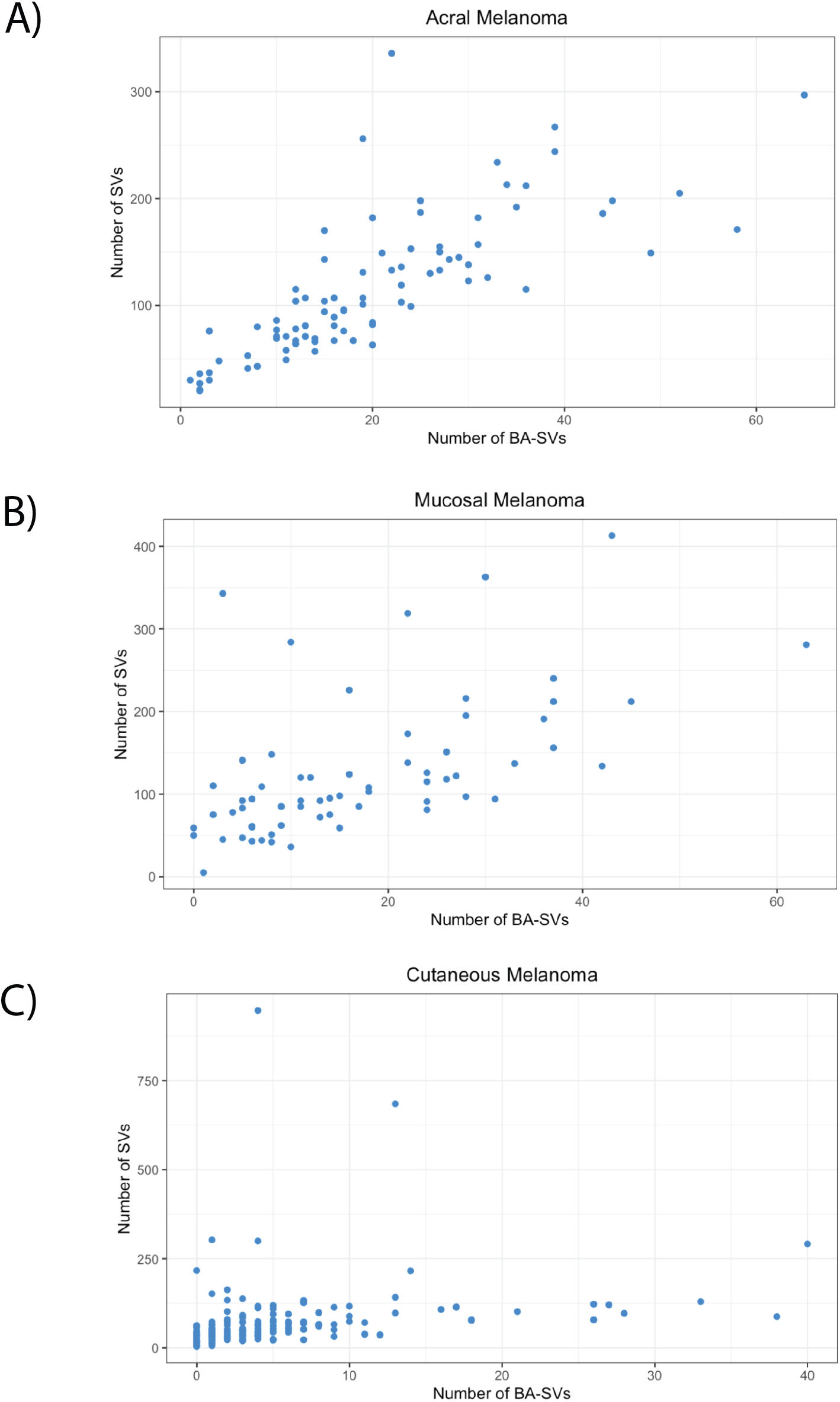
SV burden associations with BA-SV frequency. **(a)** There is a strong association between SV burden and BA-SV frequency in acral melanomas (Pearson’s r = 0.75, p = 2.2 × 10^-16^), compared to a **(b)** moderate association in mucosal melanomas (Pearson’s r = 0.57, p = 8.8 × 10^-7^), and a **(c)** weak association in cutaneous melanomas (Pearson’s r = 0.26, p = 6.2 × 10^-4^).

**Supplementary Figure 3:**
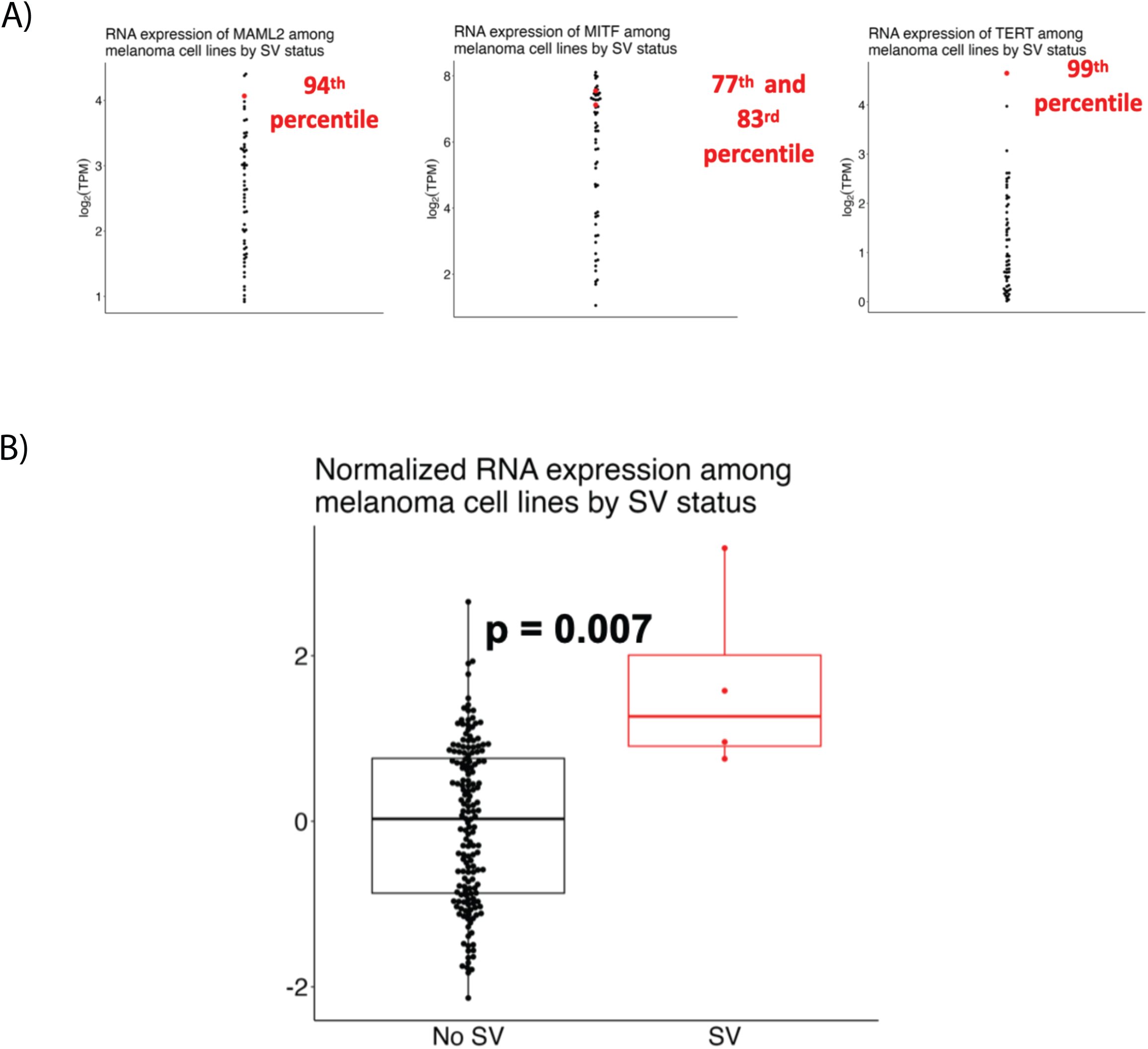
Functional consequences of highlighted and overlapping SVs in CCLE. **(a)** Percentile gene expressions of cell lines with SVs affecting genes highlighted in Figure 3. **(b)** Boxplot showing the normalized expression of genes affected by SVs compared to those that are unaffected.The p-value is from performing a Mann-Whitney test on the pan-gene aggregated z-scored (i.e. each gene z-scored separately to put the expression values on the same scale) expression.

**Supplementary Figure 4:**
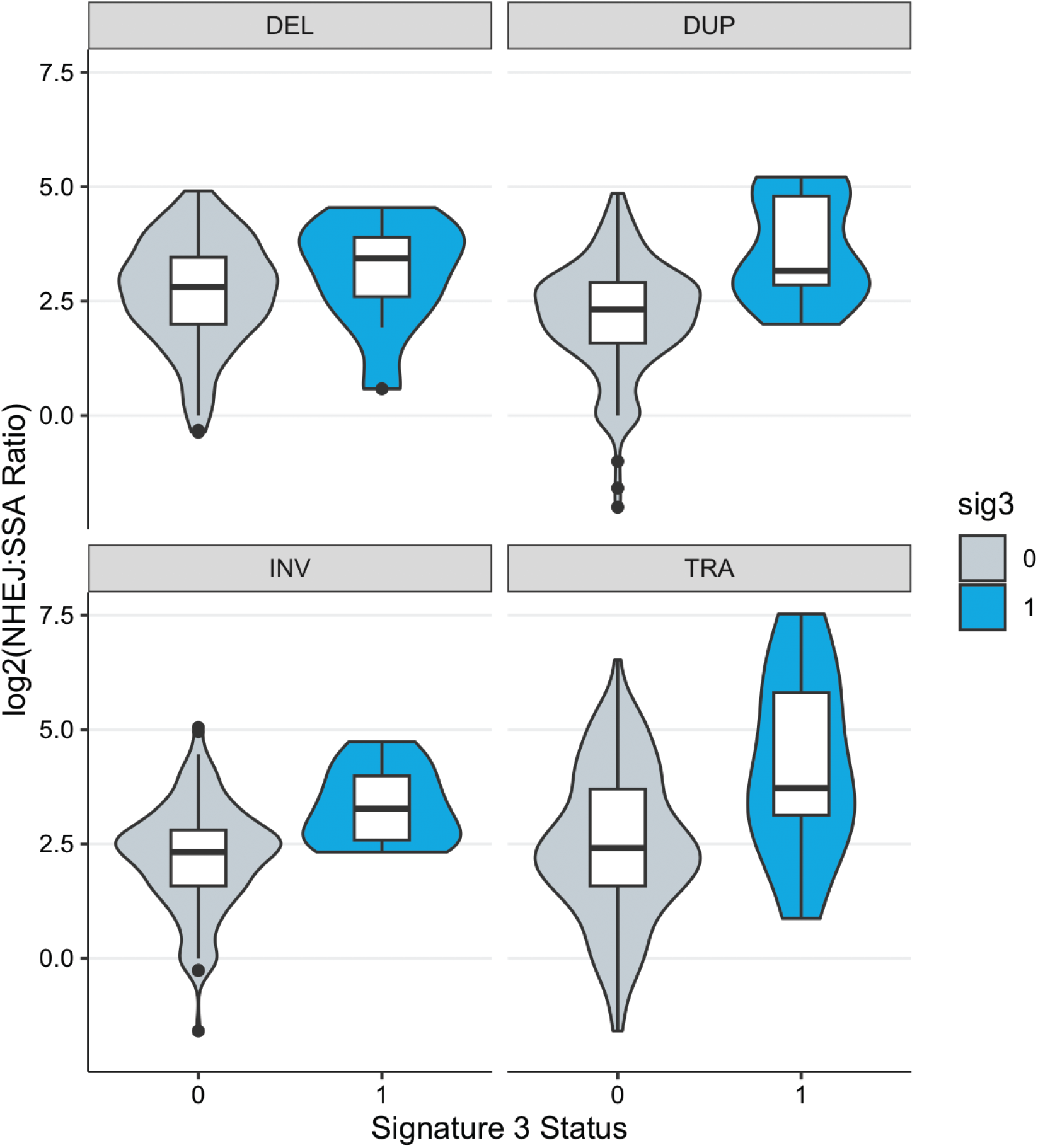
Distribution of NHEJ:SSA ratio between signature 3 and non-signature 3 tumors by SV type. The difference between the NHEJ:SSA ratio in SBS3-positive and negative tumors was statistically significant and equally as enriched in SBS3 positive tumors across DUP, INV, and TRA events (Mann-Whitney U, p < 6.9 × 10^-3^), but not DEL events (Mann-Whitney U, p > 0.05).

**Supplementary Figure 5:**
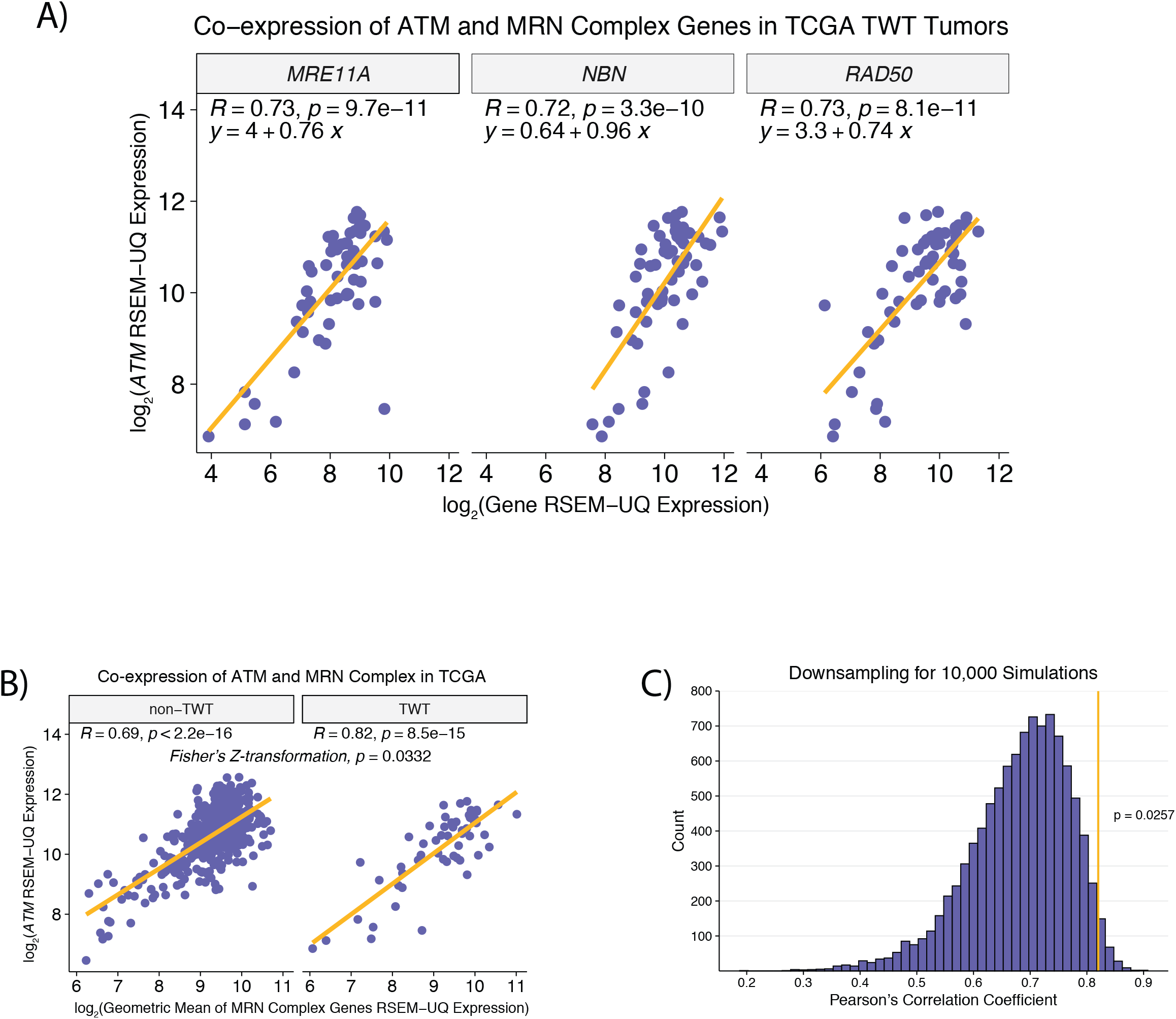
Association between *ATM* and the MRN complex in TWT and non-TWT Melanomas. **(a)** No difference was observed in the association between *MRE11A* or *NBN* expression and *ATM* expression compared to the association between *RAD50* and *ATM* expression in TWT tumors **(b)** The correlation between *ATM* expression and MRN complex expression (methods) in non-TWT and TWT cutaneous melanoma tumors. **(c)** The distribution of Pearson’s correlation coefficients from 10,000 randomly sampled simulations where non-TWT cutaneous tumors are downsampled to the number of TWT cutaneous tumors in the cohort.

**Supplementary Figure 6:**
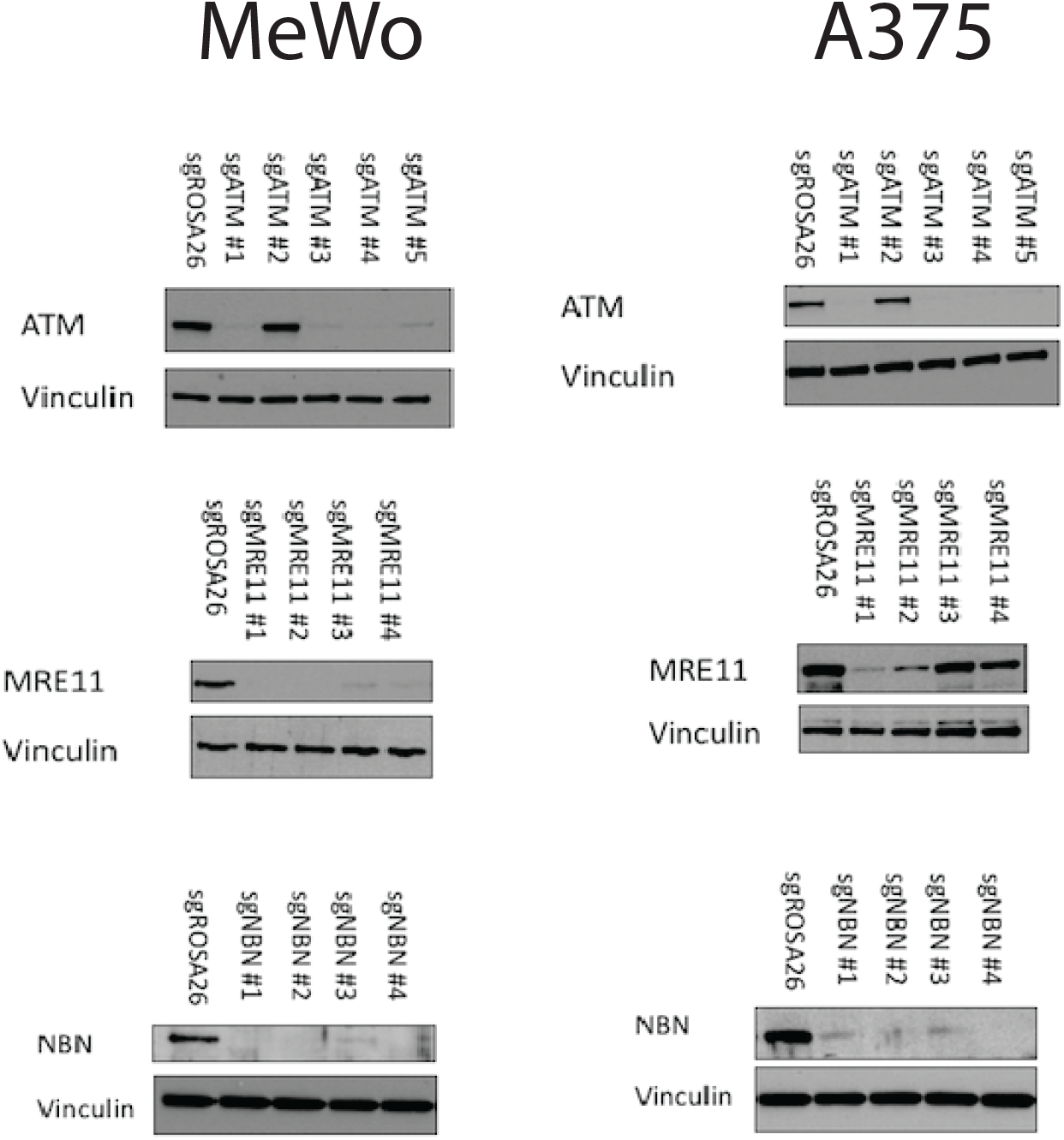
Western Blots for MeWo and A375 melanoma cell lines. Western blots for the MeWo and A375 melanoma cell lines for knockouts of *ATM*, *MRE11*, and *NBN*. Vinculin is used a negative control.

